# Synanthropic Culicidae (Diptera) in three fragments of Atlantic Forest in northern Parana, Brazil

**DOI:** 10.1101/2023.10.31.565019

**Authors:** Luiz Eduardo Grossi, Leticia Bernardete da Silva, Vinicius Martins Novaes, Halison Correia Golias, Mario Antonio Navarro da Silva, João Antonio Cyrino Zequi

## Abstract

Synanthropic Culicidae were sampled using natural and artificial traps in three forest fragments (Daher Wood, Botanical Garden, and Mata dos Godoy State Park) in Londrina, Paraná, Brazil. To collect Culicidae, six traps were distributed in two separate transects of 70 m each, parallel, with a distance between them of 30 m. Three traps were natural (bamboo) and the other three were artificial (tires). In total, we collected 15,048 specimens distributed in eight species. The peri-urban fragments showed the highest richness. Tires were preferred as breeding sites. The seasons of highest and lowest sampling were summer and winter. The Daher Wood and Botanical Garden showed high similarity, and the Godoy Wood, an intact environment, showed the greatest dominance. The mosquitoes demonstrated varying degrees of synanthropy. Correlations were positive between temperature (0.53) and precipitation (0.40) with Culicid abundance. The Principal Component Analysis indicated that tires were most influenced by temperature, relative humidity, and bamboos by Total Dissolved Solids. Four of the collected species showed potential to be vectors of etiological agents. Abiotic factors directly influence the biology of mosquitoes, which is reflected in higher diversity in warmer and rainy periods. Urban and peri-urban areas showed more synanthropic mosquitoes due to their preference for modified sites. Vector species in these areas are of concern because they can modify disease cycles, and for this reason, it is essential to monitor these areas.

## Introduction

Urbanization and intensified population growth have changed the environment rapidly, arising from the search for housing, work, industries, commerce, and transport routes. These needs combined with poor planning, deforestation, and overexploitation of natural resources have caused major environmental impacts (De Souza & Hayashi, 2013). The urbanization process has led to natural biomes being subjected to continuous devastation, putting several ecological processes at risk, and affecting many organisms (Rezende et al., 2018).

Tropical forests are being fragmented around the world (Arroyo-Rodrigues et al., 2017). Forest fragmentation causes the loss of species in modified environments (Myers et al., 2000; Fahrig, 2003). This process, caused by the destruction of micro-habitats, alters the patterns of dispersal, migration, distribution, behavior, and survival of species (Laurance, 2008; Ferreira, 2019).

The Brazilian Atlantic Forest originally occupied an area of approximately 1.5 million km^2^, and today is reduced to 28% of its original coverage (Rezende et al., 2018). Currently, the Atlantic Forest is considered one of the most threatened tropical forests on the planet (Safar; Magnago; Schaefer, 2020), since more than 60% of the Brazilian population resides in this biome (Scarano; Ceotto, 2015).

The imbalance caused by the fragmentation of biomes can favor populations of species that are more adapted and resistant to environmental alterations. Some species of mosquitoes present great genetic and ecological plasticity (Zequi et al., 2005, Duarte et al., 2022; Fimia et al., 2022) having a high capacity to adapt to the anthropic environment, responding better to environmental pressures, and becoming abundant in areas that now have low richness (Chaves et al., 2011; Guedes, 2012; Nascimento et al., 2021). Urban fragments that suffer human action without adequate monitoring for this group, may contain, in addition to wild species, opportunistic, synanthropic, and vector species, as these locations contain natural and artificial breeding grounds used for oviposition (Zequi et al., 2005). In this context, synanthropic culicid larvae tend to occupy specific types of breeding sites (Da Silva, 2002).

Several characteristics of breeding sites influence a pregnant female when choosing a location for oviposition, such as color, consistency, size, and shape, among others (Lopes et al., 1995; Nunes-Silva et al., 2020). When a female chooses an artificial container to lay her eggs, it may just be a type of opportunism, but it may also be a case of a change in her habits (Lopes, 1997; Almeida et al., 2020; Evangelista et al., 2021). In altered landscapes, changes occur in the distribution and supply of breeding sites, favoring synanthropic species (Montagner, 2014; Almeida et al., 2020).

Changes in the behavior of mosquito vectors may have epidemiological potential (Sereno, 2022), enabling the emergence or reappearance of new diseases (Marques, 2014; Bellini et al., 2022). The change in population dynamics caused by human actions often benefits these vector species, impacting human populations that live nearby or that carry out activities nearby (Hutchings et al., 2005; Guedes & Navarro-Silva, 2014).

Although studies with culicidae have been performed in the Brazilian Atlantic Forest in the state of Paraná (Santos et al., 2019; Da Silva et al., 2019), most of them are concentrated in the coastal and southern regions of the state. Therefore, the current work aimed to study the occurrence of synanthropic culicidae species in an urban, a peri-urban (anthropized), and a preserved forest fragment of the Brazilian Atlantic Forest. We sought to evaluate the diversity of these species in different areas, verify preferences regarding artificial and natural breeding sites for oviposition, evaluate the degree of synanthropy, observe seasonal fluctuations, and distinguish vector species from etiological agents for monitoring and control purposes.

## Methods

The study area is in the municipality of Londrina, north of Paraná, Brazil, with geographic coordinates 23°08’47” and 23°55’46” south latitude and 50°52’23” and 51°19’11” west longitude. The municipality has a territorial area of 1,652.569 km² and an estimated population of 555,937 inhabitants. The demographic density corresponds to 336.41 inhabitants/km², with a Human Development Index considered high 0.778 (IBGE,2023).

The climate in Londrina is characterized as hot subtropical, with a long, muggy summer and cloudy skies and a short winter. Throughout the year the temperature varies between 13 °C and 30 °C, with precipitation. The hot season lasts 6.1 months, from October to April, with an average daily temperature of 28°C, and January is the hottest month, with a maximum of 30°C and a minimum of 21°C, on average (Weatherspark, 2023).

The season with the highest rainfall lasts 5.4 months, from October to March. January is the month with the highest number of days of rainfall, with an average of 18.4 millimeters of rainfall (Weatherspark, 2023).

Sampling was carried out in three forest fragments of the Brazilian Atlantic Forest, located in the municipality of Londrina, Daher Wood (MD) (23° 18’ 55” S and 51° 12’ 16” W), a peri-urban fragment, the Botanical Garden (BG) (23° 21’ 44” S and 51° 10’ 22” W), and a preserved fragment in the Mata dos Godoy State Park (MG) (23° 26’ 53” S and 51° 15’ 21” W) (Figure 1) between July 2016 and May 2017, covering the four seasons of the year.

**FIGURE 1:**
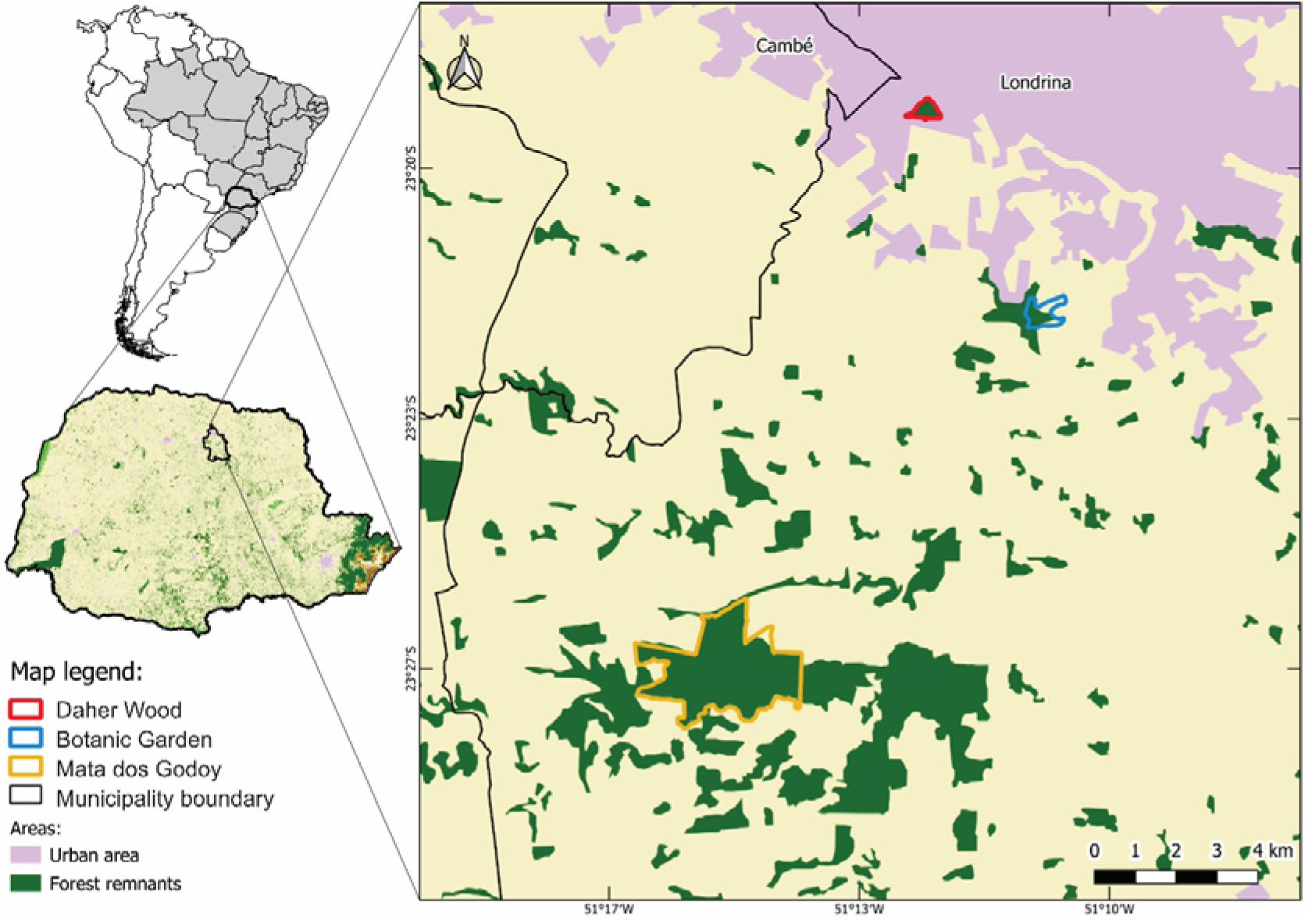
Forest fragment areas of the Atlantic Forest sampled in the municipality of Londrina, North of Paraná, South of Brazil, between July 2016 and May 2017. Source: SOS MATA ATLANTICA FOUNDATION AND NATIONAL INSTITUTE FOR SPACE RESEARCH – INPE, 2018.

Daher Wood is an urban forest reserve that occupies an area of 3.3 hectares, located in the urban area of Londrina. It is adjacent to the Celso Garcia Cid highway, PR 445, km 380, and contains a large variety of native plant species. The lower stratum is composed of secondary vegetation with an abundance of lianas and monocotyledons of the genus *Merostachys* Spreng, 1824.

The Botanical Garden is a peri-urban area, with more than 100 hectares of native forest, exotic species, springs, and rivers.

The Mata dos Godoy State Park (MG) is home to several species of fauna and flora threatened with extinction and is one of the last native forest reserves in northern Paraná. It contains 790 hectares of Semideciduous Seasonal Forest. Approximately half of the Park is a flat area (600 m a.s.l^1^), and the remainder being inclined at 470 m, reaching the lowest portion at the southern limit, where Ribeirão dos Apertados is located.

Two transects of 70 m were designated in each fragment, in parallel, with 30 meters between them, and containing three traps each, one fragment contained natural traps and the other artificial traps. For the artificial traps, car tires were used – 13-inch rims, prepared, cut, and placed in the designated locations. For the natural traps, bamboo internodes (*Bambusa* sp.) were used, measuring 25 cm high and 12.5 cm wide, called internode traps. In each transect, the traps were positioned 10 m from the edge of the fragment with a spacing of 30 m between them, at 10 m, 40 m and 70 m away from the edge, all at ground level. After being positioned, distilled water was added to the traps (tire: 1.5 liters and bamboo: 1 liter) (Figure 2).

**Fig. 2.**
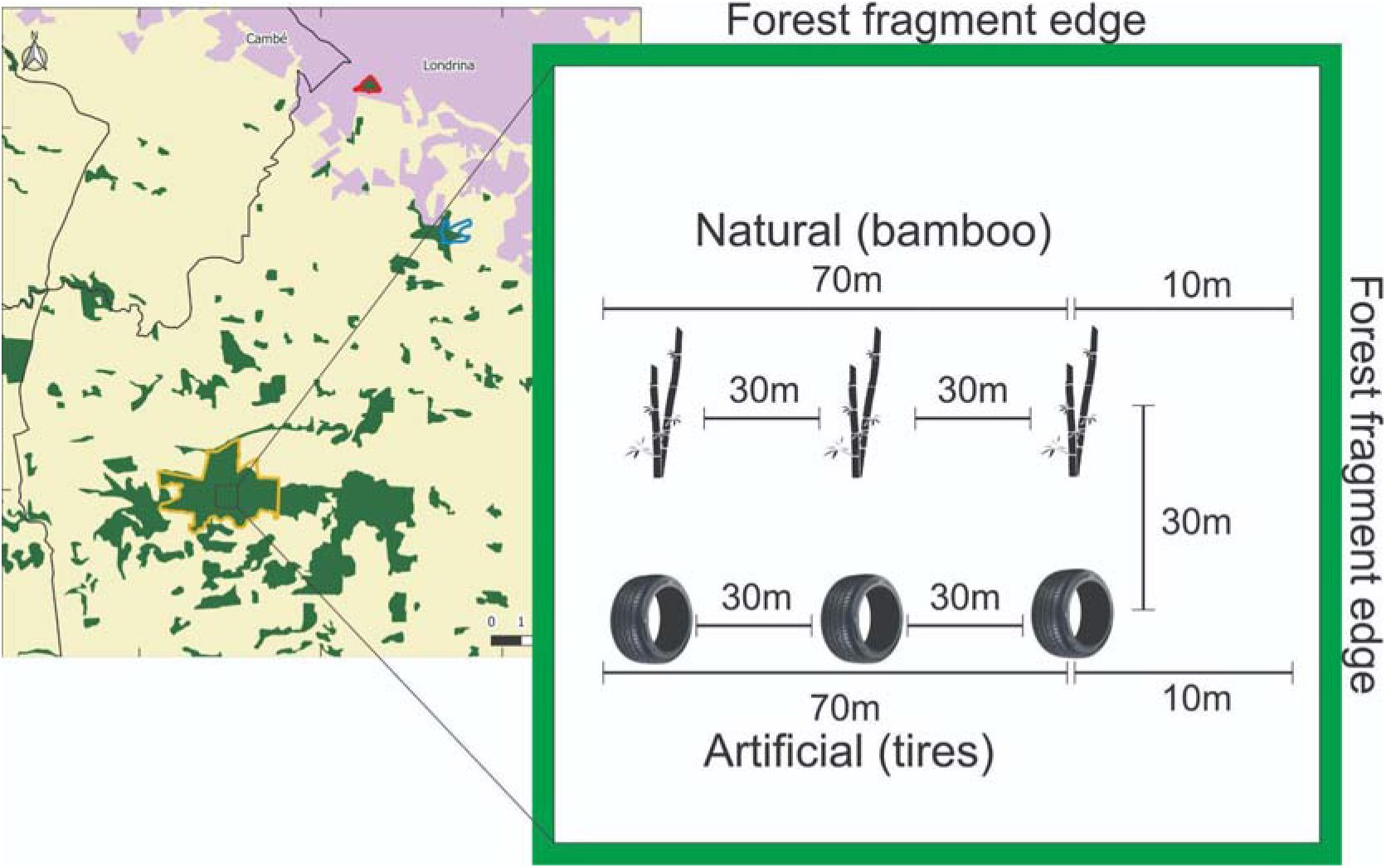
Schematic representation of data collection carried out in three forest fragments of the Brazilian Atlantic Forest located in the Northern Region of Paraná, Brazil between 2016 and 2017. (The figure is schematic and not to scale).

The total volume of water in the traps was replenished weekly during the collection periods. Physicochemical parameters of the water (pH, temperature, conductivity, salinity, dissolved and saturated oxygen, and TDS) from the breeding sites were investigated with the aid of a Hanna^Ⓡ^ multiparameter, model HI 9828. The temperature and relative humidity of the air were continuously monitored with the aid of a thermos-hygrometer installed in the fragments throughout the sampling period at each station, positioned between the transects. Precipitation data were provided by the Agronomic Institute of Paraná (IAPAR) (23° 22’ S and 51° 10’ W). Sampling was carried out quarterly. In each quarter, five consecutive weekly collections were performed per fragment, including abiotic data collections. To capture the immature Culicidae, the water from the breeding sites was strained using a sieve (1 mm mesh; 12 cm in diameter) and the Culicidae were placed in plastic bottles with ventilated lids (7.5 cm in diameter and 400 mL). Finally, the water volume of the breeding site was restored, and the larvae and pupae were transferred to the Medical Entomology Laboratory at the State University of Londrina for breeding and subsequent screening. The collected specimens were deposited in the Entomological Collection of the State University of Londrina.

Immature Culicidae with low abundance and with real difficulty in identification based on larval morphology were raised until they reached the adult stage, thus obtaining the exuviae of the fourth instar for comparisons in identification. The rest of the larvae were raised to the fourth instar and then mounted on semi-permanent slides with Hoyer’s liquid. To identify the specimens, the dichotomous keys of Consoli & Oliveira (1994), Forattini (2002) and WRBU (2018) were used with the aid of an Olympus^Ⓡ^ CH30LF100 stereo optical microscope. The identified species were confirmed with specimens from the Padre Jesus Santiago Moure Entomological Collection at the Federal University of Paraná.

To calculate the degree of synanthropy of the Culicidae, the Nuortueva (1963) index was used, with modification. The index uses the following formula: IS = 2ª + b – 2c/2, where: a = percentage of a species x sampled in the urban area in relation to the same species sampled in the rural area and in the forest; b = percentage of species x in rural areas; c = percentage of species x in the forest. In this study, the areas were altered, as follows: a = fragment within the municipality (Daher); b = fragment located in the peri-urban area (Botanical Garden); c = fragment of forest (Godoy). The index has values ranging from + 100 to – 100, with positive values revealing a high degree of synanthropy and negative values showing the opposite.

To measure diversity, the Shannon-Wiener (H’) and Margalef (Dmg) indices were used; for species dominance, the Simpson (C) and Berger-Parker (d) indices were used; and to verify equitability the Shannon-Wiener (eH’) (e = H’/ log S) and Pielou (J) indices were chosen, performed using the DiVes Program – Species Diversity 3.0 (Rodrigues, 2014). The similarity between the fragments was calculated and expressed by the Morisita and Bray-Curtis Index, performed in Past 3.0 Software (Hammer, 1999). To test differences in the richness and abundance of mosquitoes between fragments and between types of breeding sites, the T-test, ANOVA, and Kruskal-Wallis tests were used with Fisher’s post-hoc for the former and Dunn’s test for the latter in Statistica Software (Statistica 7.0, 2005).

Regarding sampling, species rarefaction curves were created for the fragments and species richness was estimated with the help of the non-parametric Jackknife (I and II) and Chao (I and II) extrapolator indices (estimators). The data used to assemble the curves and calculate estimators were obtained through a randomization process in the Estimate-S 9.1.0 program (Colwell, 2013). Temperature, relative humidity, and precipitation data were related to insect abundance using linear correlation with Statistica Software (Statsoft 7.0). To verify which abiotic variables most influenced the two types of breeding sites, Principal Component Analysis (PCA) was used in Past 3.0 Software. (Hammer, 1999).

## Results

During the sampling period, 15,048 specimens of Culicidae were collected, distributed across five genera and eight species. The peri-urban fragment (BG) presented the greatest richness, with eight species, followed by the urban fragment (MD), with seven species, and finally the preserved environment (MG), with five species. The most abundant species were *Cx. eduardoi* and *Li. durhamii* and the least abundant were *Cx. saltanensis* and *Ae. aegypti* (table 1).

**TABLE 1:**
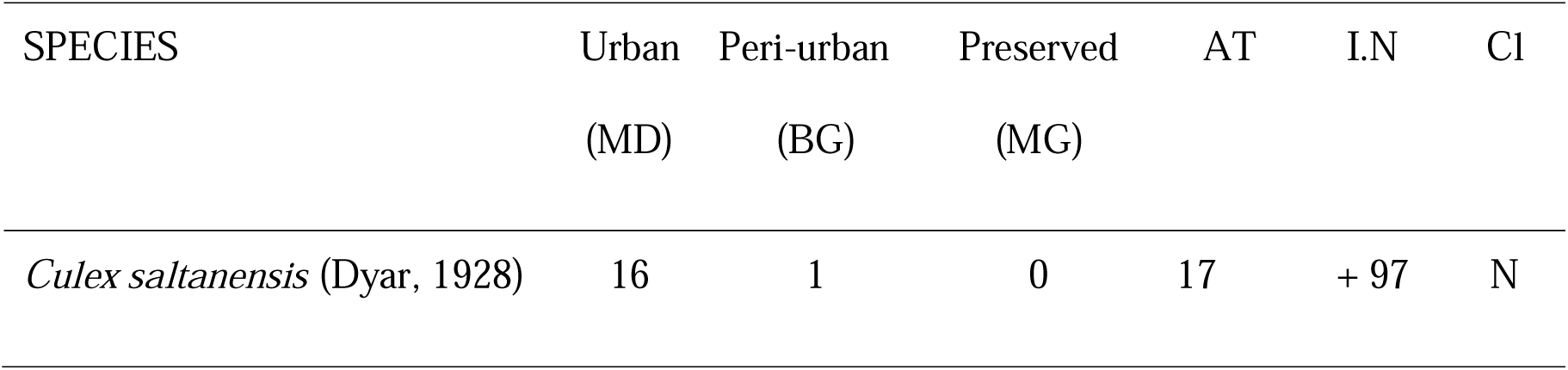
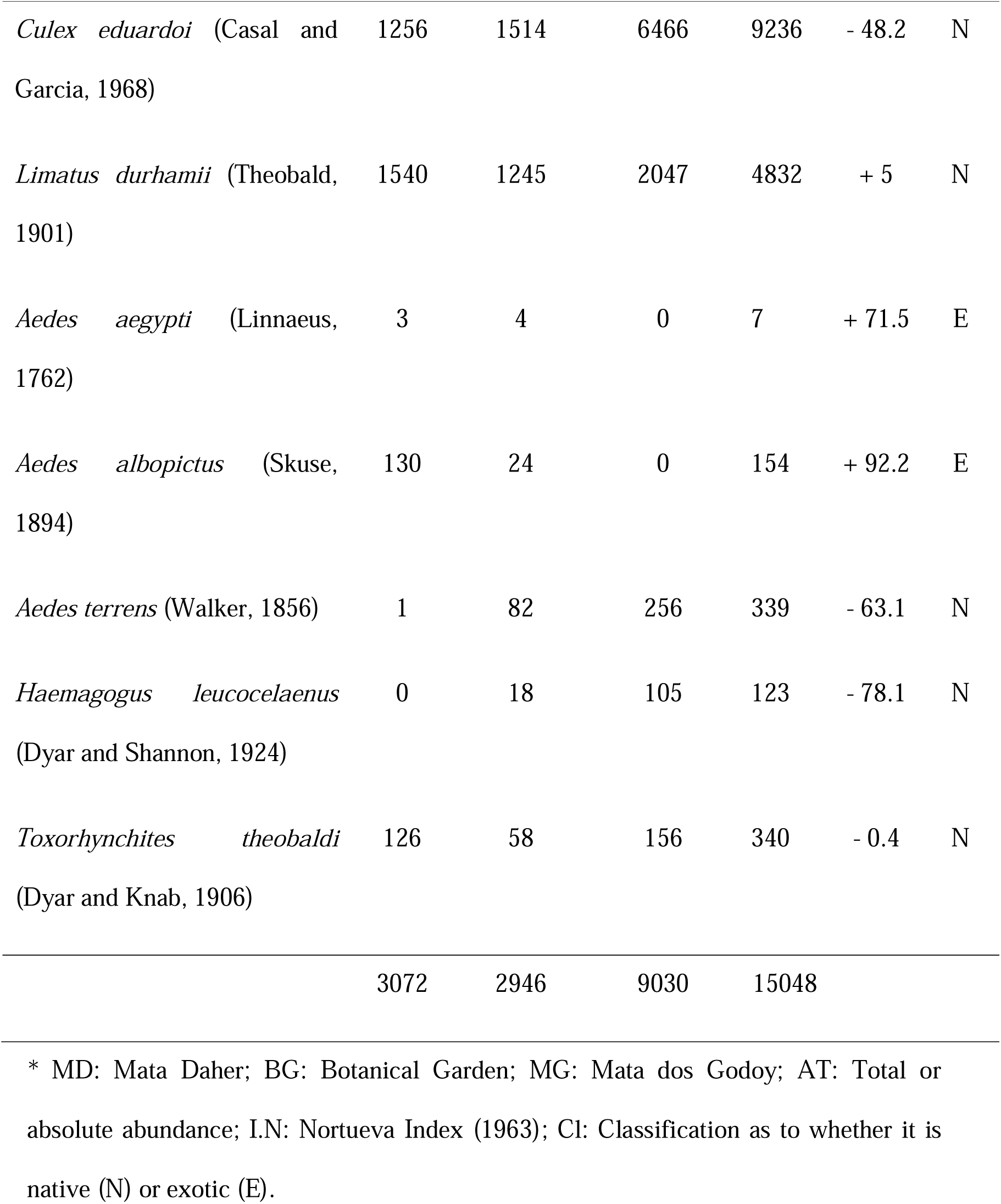
Total abundance, Nortueva Index, and classification (native or exotic) of Culicidae species collected in forest fragments of the Brazilian Atlantic Forest between July 2016 and May 2017 in the Northern region of Paraná, Brazil.

*Culex eduardoi* was the most abundant species in the peri-urban fragment and in the preserved area. The species *Culex saltanensis*, *Ae. aegypti,* and *Ae. albopictus* were not sampled in the preserved fragment. Table 1 lists, quantifies, and demonstrates the degree of synanthropy of the species collected in the three fragments.

Considering three traps of each type in each area, multiplied by the number of collections, there was a total of 180 artificial traps (tires) and 180 natural traps (bamboos), that is, 60 samples of each type for each area. Artificial traps were characterized as the preferred breeding site for the mosquitoes, with a colonization rate of 90.55% and a portion of 88% of sampled larvae compared to natural breeding sites, which had a colonization rate of 38.89% and a portion of 12% of larvae found. Furthermore, artificial breeding sites also showed greater productivity when considering population density and species diversity, as seven species were found mainly in artificial breeding sites.

The Nortueva index (1963) revealed different degrees of synanthropy for the culicidae sampled (Table 1). These values are related to the habits of each species, whether it is found with greater prevalence in artificial habitats.

The genus *Aedes* was represented by three species. *Aedes aegypti* was found in both breeding sites, with a preference for artificial sites. The few representatives of this species were sampled only in autumn in the urban and peri-urban fragment. *Aedes albopictus* was more abundant during spring and summer, revealing no specific preference for breeding sites. *Aedes terrens*, despite being sampled in both breeding sites, revealed a preference for artificial breeding sites, with peak abundance in summer, and without representatives collected in winter.

*Aedes albopictus* was found in the urban and peri-urban fragments. Larvae developed both in natural microhabitats (such as bamboo internodes and tree hollows) and in a wide range of artificial containers (Lopes et al., 2002), which resulted in the same percentage of individuals in both types of breeding sites for this study. This species was found in all seasons, with greater abundances in spring and summer. Populations are able to remain at lower temperatures (Lima-Camara, 2010), but decline as rainfall decreases (Leandro, 2012).

*Aedes terrens* was collected in the three areas, but with most individuals in the preserved area and with a preference for oviposition in artificial breeding sites. This species was not found cohabiting with *Ae. aegypti* and *Ae. albopictus,* revealing differences in the choice of colonized environments. However, it is often found cohabiting with most species that use tires as breeding sites in their habitat (Lopes, 1997). The greatest abundance occurred in summer, with this species preferring a high temperature and precipitation. High population averages in hot and rainy months were also observed by Zequi et al. (2005).

*Culex saltanensis* and *Ae. aegypti* had few representatives, due to the fact that they are species found closer to human populations. *Culex saltanensis* is commonly found in treatment ponds with large amounts of organic matter (Zequi et al., 2005) and *Aedes aegypti* is highly adapted to the urban environment (Johnson et al., 2002) with highly synanthropic and anthropophilic behavior (Natal, 2002).

*Haemagogus leucocelaenus* showed a preference for artificial breeding sites, demonstrating that this species has the ecological valence to colonize this type of breeding site (Lopes, 1997; Zequi et al., 2005). The peak abundance of *Hg. leucocelaenus* larvae was in the summer and they were not found in seasons with lower temperatures and little precipitation. Peaks of abundance occur in warmer periods (Zequi et al., 2005). Despite being found in anthropic environments, *Hg. leucocelaenus* is better adapted to non-urban environments (Montagner et al., 2017). The frequent cohabitation of this species with *Ae. terrens* highlights the wilder habits of both (Zequi et al., 2005).

*Toxorhynchites theobaldi* presented its peak abundance in the summer, with fewer individuals in the winter. They were found cohabiting with all the other species sampled, an important fact, since the larvae of this genus are predators and feed on other larvae in the breeding site (1). In 75% of positive traps for this species, the number of larvae was low, ranging from one to three. Either isolated egg laying helped prevent cannibalism, or cannibalism itself caused this number to be low (Lopes, 1997). *Tx theobaldi* was found in all areas, revealing its adaptive capacity in anthropic environments (Montagner et al., 2017).

Of the eight species sampled, only four were collected in winter, totaling 696 individuals. In the spring, five species and 2755 individuals were collected, in the summer, six species and 7295 individuals, and in the autumn, eight species were collected, totaling 4302 individuals (Figure 3).

**FIGURE 3:**
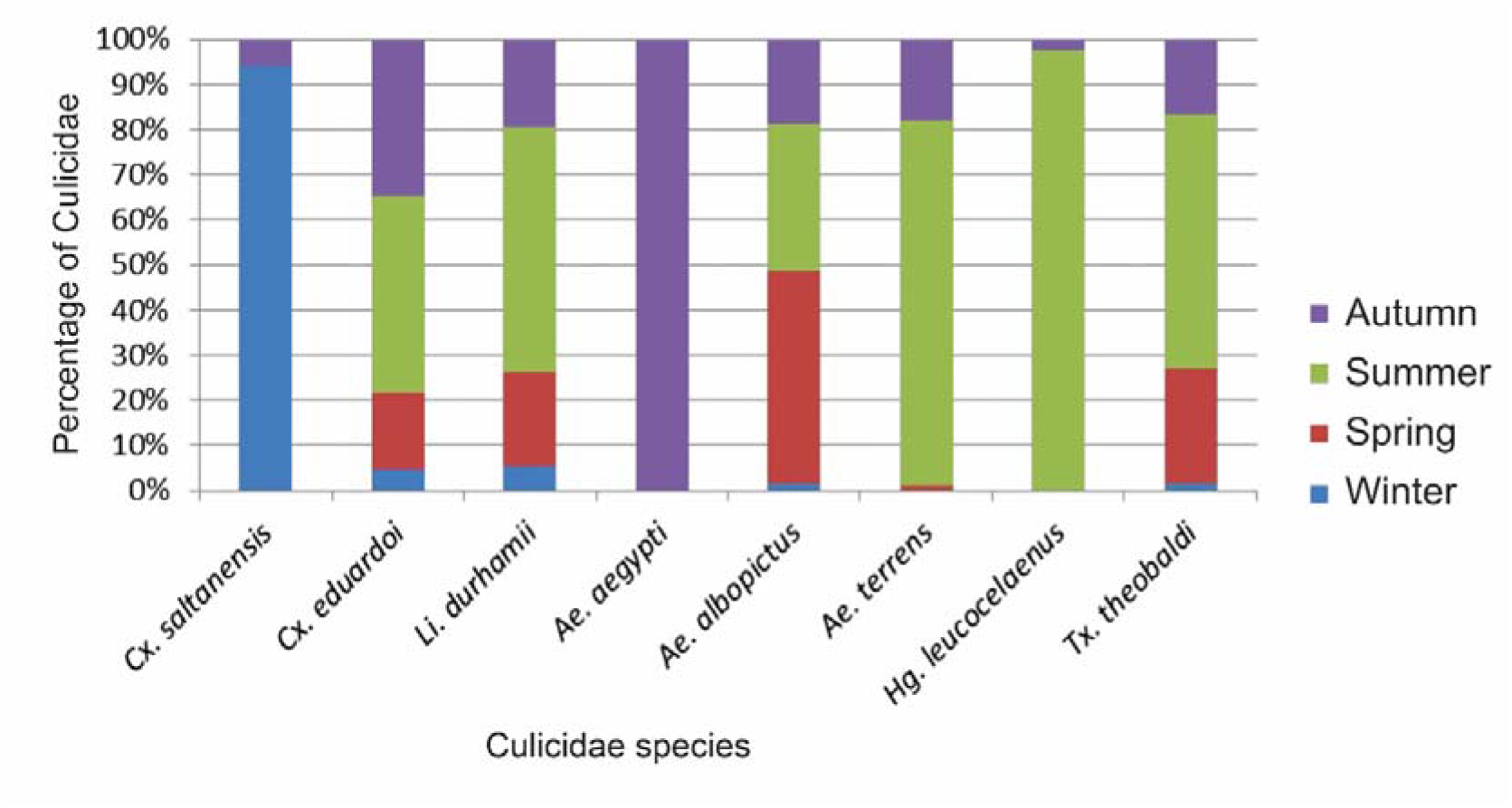
Percentage of Culicidae species sampled per season in the three forest fragments of the Brazilian Atlantic Forest between July 2016 and May 2017, in the Northern region of Paraná, Brazil.

The urban and peri-urban fragments were similar in terms of diversity (table 2). The Morisita index showed similarity values between 0.976 and 0.992 for these two fragments, and the Bray-Curts index values between 0.84 and 0.9.

**TABLE 2:** Diversity, Dominance, and Evenness Indices of Culicidae collected in three forest fragments of the Brazilian Atlantic Forest between July 2016 and May 2017, in the Northern region of Paraná, Brazil.

| Areas | Diversity |  | Dominance |  | Evenness |  |
| --- | --- | --- | --- | --- | --- | --- |
|  | Shannon | Margalef | Simpson | B. Parker | Shannon | Pielou |
| Urban (MD) | 0.4401* | 1.7205 | 0.4218 | 0.5013 | 0.2218 | 0.5208 |
| Peri-urban (BG) | 0.4192 | 2.0177* | 0.4438 | 0.5139 | 0.1901* | 0.4641* |
| Preserved (MG) | 0.3468 | 1.0112 | 0.5653* | 0.7161* | 0.2829 | 0.4962 |

\* Reveals which values were most representative.

The greatest dominance was found in the preserved fragment due to the low richness of the fragment and the high abundance of two species, *Cx. eduardoi* and *Limatus durhamii*. The peri-urban fragment was the area with the best evenness due to its greater richness and a more homogeneous distribution of species (Table 2).

Regarding abundance, there was a significant difference only between the peri-urban and preserved fragments (p= 0.015). Types of breeding sites in total were also significantly different with (p= 0.000001), as well as types of breeding sites by area, with (p= 0.000012) in the urban fragment, (p= 0.00026) in the peri-urban fragment, and (p = 0.000008) in the preserved fragment. When comparing natural breeding sites between the three areas, there were significant differences between the urban and preserved fragment (p= 0.0103) and between peri-urban and preserved fragment (p= 0.006). Artificial breeding sites (tires) did not show significant differences between areas. Regarding species richness, there were no significant differences in any of the cases compared.

Sampling during all climatic seasons made it possible to discover a greater number of species, as they have preferences for certain periods, influenced by environmental characteristics and the biology of each species. Regarding mosquito abundance, statistical data showed a moderate correlation of 0.53 with temperature and a correlation of 0.40 with precipitation. Relative air humidity did not show a significant correlation. Figure 4 presents the abundance of culicidae related to abiotic factors.

**FIGURE 4:**
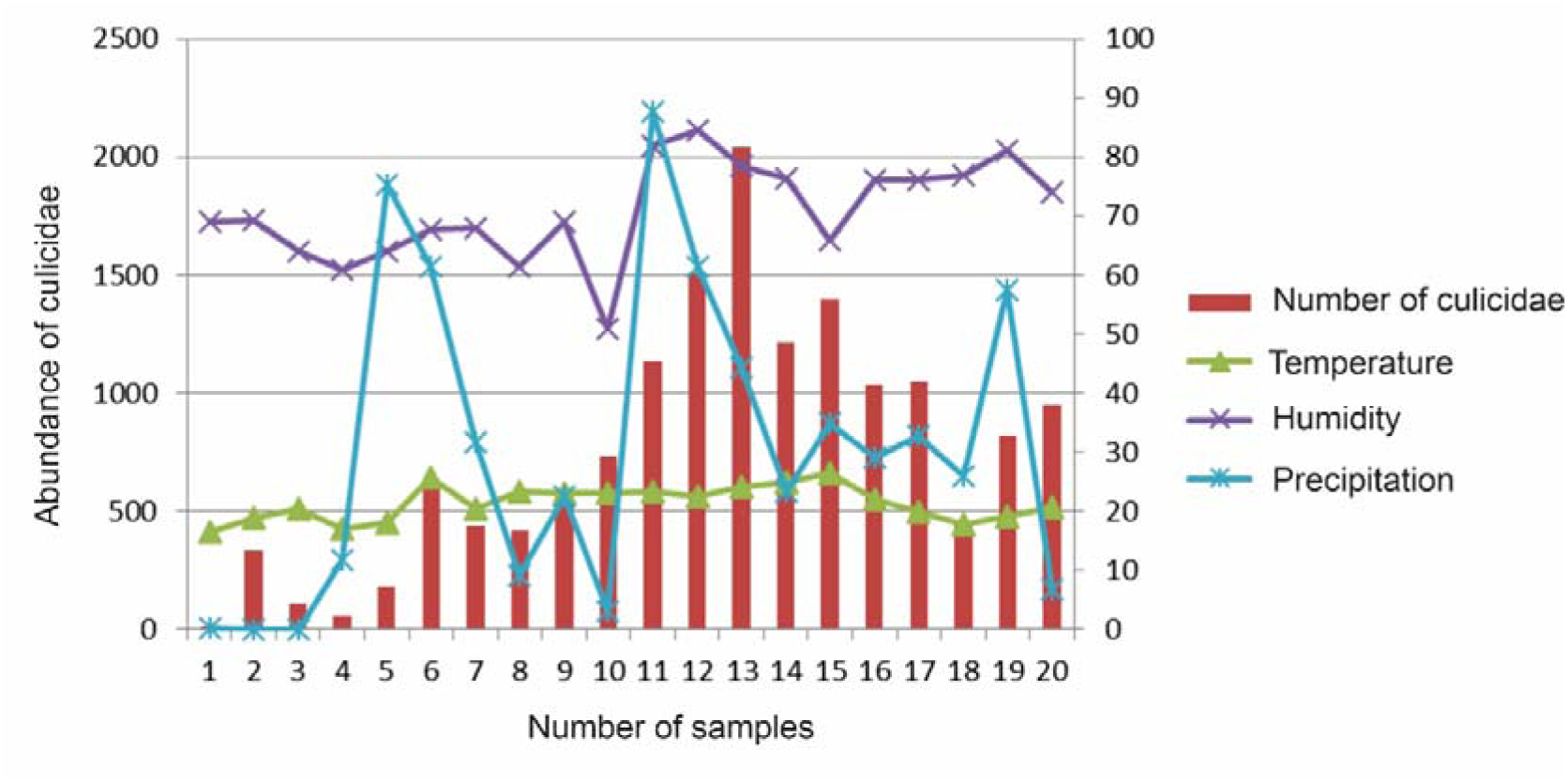
Abundance of Culicidae collected between July 2016 and May 2017 in three forest fragments of the Brazilian Atlantic Forest and their relationship with temperature, relative air humidity, and precipitation in the Northern region of Paraná, Brazil.

The PCA (Appendix – Figure 17) revealed that the tires are close to each other as well as the bamboos to each other, making it possible that the area is not influencing the choice of breeding site. The artificial breeding site was more influenced by relative air humidity and precipitation (external to the breeding site) when compared to other variables in the internal area of the breeding sites (pH, salinity, water temperature, conductivity, TDS (Total Dissolved Solids), dissolved oxygen, and % oxygen. The natural breeding site, on the other hand, did not obtain such an explicit approximation of the abiotic variables, being only slightly closer to the TDS variable, when compared to other variables. This result shows that environmental factors in the fragment have greater influence when compared to factors in the breeding site itself. The low colonization of natural breeding sites is associated with the biology of the mosquitoes that chose the artificial breeding site, however the PCA allowed us to verify that the amount of organic matter in these breeding sites was important, confirming the greater colonization by *Cx. eduardoi*, which is prevalent in places with lots of organic matter (Lopes et al., 2012).

The low richness obtained by the Chao I, Chao II, Jackknife I, and Jackknife II estimators shows that despite estimating a few more species for the fragments, this number was not significant (Table 3). Figure 5 presents the species rarefaction curves for the three fragments, where the sampling sufficiency can be observed.

**FIGURE 5:**
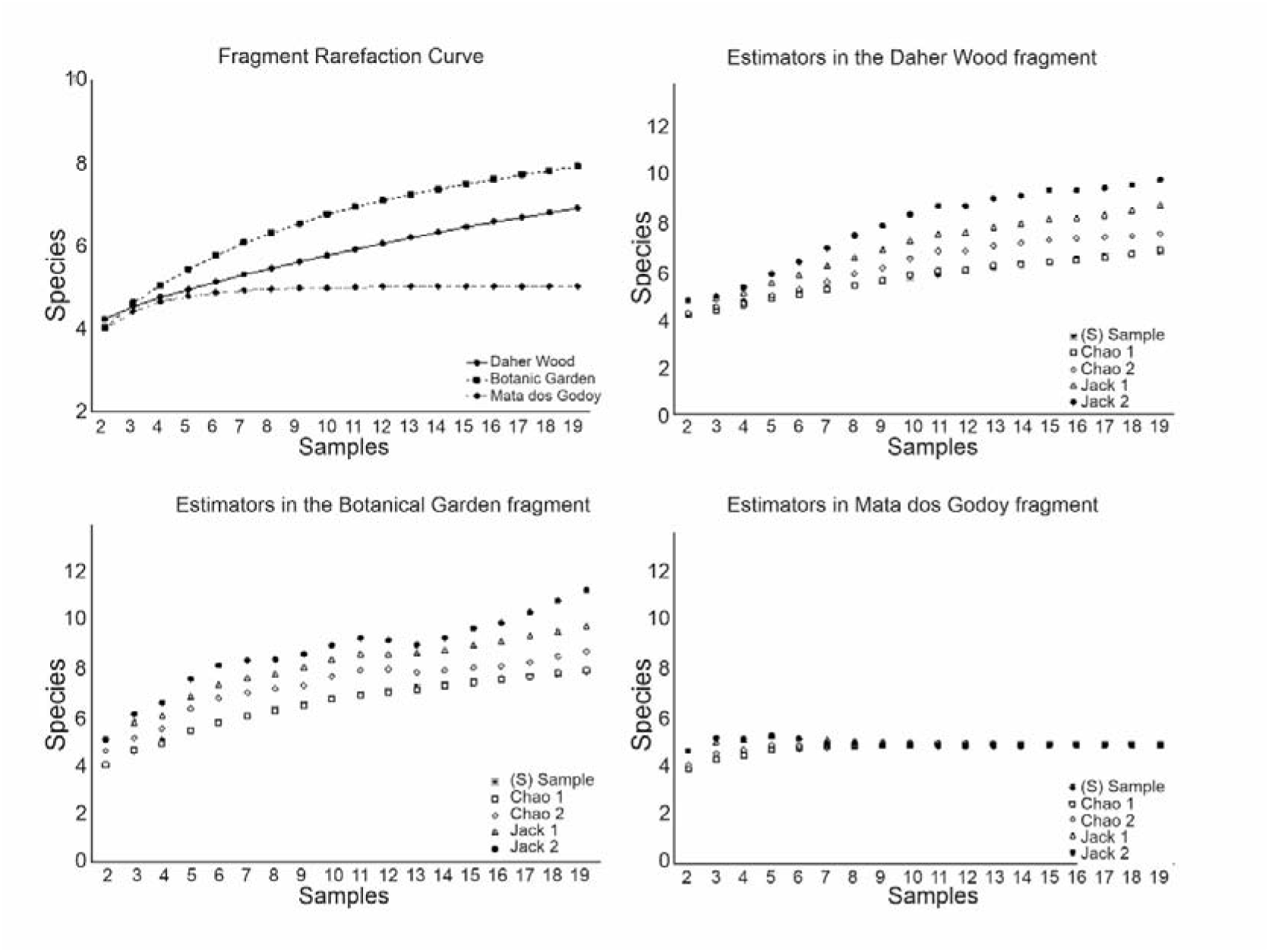
Rarefaction curves of Culicidae species sampled in the three fragments of Brazilian Atlantic Forest evaluated in the period between July 2016 and May 2017

**TABLE 3:** Chao I, Chao II, Jackknife I, and Jackknife II estimators, showing the number of species estimated for each fragment studied.

|  | Chao 1 | Chao 2 | Jackknife 1 | Jackknife 2 |
| --- | --- | --- | --- | --- |
| Daher | 7 | 7.48 | 8.9 | 9.93 |
| Botanic Garden | 8 | 8.95 | 9.9 | 11.7 |
| Godoy | 5 | 5 | 5.39 | 9.95 |

## Discussion

In fragmented habitats, there is a loss of species richness, with the extinction or migration of wild species (Dos Anjos & Navarro-Silva, 2008). However, some Culicidae species are able to adapt to new selective pressures in this altered environment (Forattini & Massad, 1998; Chaves et al., 2011). The greater richness in urban and peri-urban fragments revealed that the species sampled prefer altered locations. In these environments there is strong anthropic action, with the disposal of garbage, tires, and other containers, that increase the number of artificial breeding sites, and which end up becoming refuges for species of mosquitoes with a synanthropic nature (Zequi et al., 2005). More preserved areas tend to have a greater richness of wild culicidae species (Dos Anjos & Navarro-Silva, 2008), however, in this case, as they are synanthropic culicidae, the situation is reversed, and because of this and other factors the preserved fragment showed the least richness.

The preference for artificial breeding sites reveals the plasticity and/or synanthropism of the species in the search for places for oviposition. They benefit from environmental changes caused by man (Gomes & Forattini, 1990) and take advantage of containers with stored water for oviposition (Zequi et al., 2005).

The species *Aedes aegypti* is associated with humans and with high dependence on manufactured containers for oviposition (Natal, 2002). The presence of *Ae. aegypti* in the peri-urban fragment reveals some situations: the presence of garbage and other discarded containers were used for reproduction of this species, which helps to keep the population low in these forest fragments; the absence of warm-blooded animals for female food; the species expanded its habitat in search of human blood and egg-laying sites; and/or its opportunistic behavior causes it to look for places for reproduction and shelter, adapting late to native environments (Montagner et al., 2017). Despite the low number of representatives sampled, *Ae. aegypti* has always been associated with *Ae. albopictus*. These are species that, despite having different niches, are distributed in the same habitats (Leandro, 2012) and develop in the same types of artificial containers. (Honório & Lourenço-de-Oliveira, 2001). The decrease in *Ae. aegypti* is related to an increase in *Ae. albopictus* (2) because in places where they co-occur, *Ae. albopictus* was shown to be a superior competitor (Leisnham et al., 2014).

The ecological valence of *Ae. albopictus* causes it to have a wide occupation, colonizing wild and anthropic environments, and natural and artificial breeding sites (Forattini, 2002; Zequi et al., 2005; Medeiros et al., 2009; Lima-Camara, 2010). However, despite its plasticity, no representatives were found in the preserved fragment, as this species has great adaptation in places of transition between urban and wild environments (Montagner et al., 2017), living up to its high degree of synanthropy.

*Aedes terrens* is a species with more wild habits, however it can be found in anthropic environments (Zequi et al., 2005). This species was known to reproduce only in natural breeding sites, such as tree holes (Neves & Faria, 1977), but began to be collected in artificial breeding sites from 1988 onwards (Zequi et al., 2005).

The occurrence of *Cx. saltanensis*, represented almost entirely in winter, showed a preference for lower temperatures and little rain, diverging from *Cx. eduardoi*. This species was shown to be adapted to milder temperatures, which are more suitable for its development and provide lower fertility losses and lower mortality rates in adult mosquitoes (rossman & Lourenço-de-Oliveira, 1996). The totality of individuals inhabiting artificial breeding sites and the majority being found in the urban environment reveal the high adaptability of this species to modified environments, in accordance with the synanthropic index. Despite this, this species can also be found colonizing natural breeding grounds (Zequi & Lopes, 2007; Zequi & Lopes, 2012).

*Culex eduardoi* and *Limatus durhamii* were the dominant species, colonizing both types of breeding sites, during all seasons in all fragments. This abundance was also verified in other studies (Lopes et al., 1995; Lopes, 1997; Zequi et al., 2005; Montagner et al., 2017). Both species were found in the three fragments, because they have euryoecious characteristics and present genetic plasticity that makes them able to survive in anthropic environments and colonize natural and artificial breeding sites (Lopes, 1997). *Limatus durhamii* could colonize different environments and breeding sites due to its high ecological valence, and its development can occur in natural and artificial breeding sites (Silva et al., 2004) being, in general, the first non-native species to establish itself (Lopes et al., 1987). *Culex eduardoi* also has a wide ecological valence and for this reason it is found in natural and anthropic environments, colonizing natural breeding grounds and artificial breeding grounds, with a preference for tires (Lopes, 1997). However, despite preferring artificial breeding sites for oviposition, these species inhabit the different areas studied and for this reason their degrees of synanthropy were relatively low. These results may be associated with the fact that *Li. durhammi* is strongly related to native forests (Montagner et al., 2017), and *Cx. eduardoi* opt for breeding sites in shady conditions and closed forests (Lopes et al., 2012), being sensitive to environmental degradation.

The population fluctuation observed for these two species indicates that both reproduced more actively in the summer, with representative populations in the winter. They are present throughout the year, but with the highest population densities in summer when temperatures are high (Lopes, 1997; Zequi et al., 2005).

The populations of culicidae species are reduced in the coldest periods and increased in the hottest and rainiest periods. This is because the temperature directly influences their development, and an increase of 1°C can cause a 54% increase in the female’s egg laying (Nascimento et al., 2022). The correlations showed that a greater abundance of mosquitoes is generally associated with higher temperatures and high precipitation rates that occur in spring and, especially, in summer, with populations being higher in the summer in the Atlantic Forest (Rona et al., 2009). However, during the sampled period the climate proved to be atypical, with low precipitation in spring, 141.5 mm, and high precipitation in autumn, with a value of 158.15 mm (IDR-Paraná, 2023), which ended up reflecting in a greater abundance in the latter.

Some relationships between larvae of different species in the breeding sites were verified. In most traps, *Tx. theobaldi* was present with *Li. durhamii* and/or *Cx. Eduardoi*, but was also found cohabiting with the other species. *Ae. aegypti* larvae were always found sharing breeding sites with *Ae. Albopictus,* and *Ae. terrens* cohabiting with *Hg. leucocelaenus*. However, *Ae. albopictus* and *Ae. terrens* did not frequently live in the same breeding place. The greatest cohabitation was in the summer, all of them occurring in the preserved fragment and always presenting the same species, namely, *Cx. eduardoi*, *Li. durhamii*, *Ae. terrens*, *Hg. Leucocelaenus,* and *Tx. theobaldi*. Due to the great abundance in the warmer seasons, the search for oviposition sites by females is greater, which is why artificial breeding sites can generally house a wide diversity of species (Zequi et al., 2005).

The preserved area has a lower number of species of synanthropic mosquitoes, which explains the very low similarity with the other fragments, according to Morisita (above 0.8) and to Bray-Curtis (below 0.5).

## Conclusion

Abiotic factors directly influence the biology of mosquitoes, which is reflected in greater abundance in hotter and rainier periods, with temperature, precipitation, and relative humidity positively affecting the diversity of this group. The preference for artificial breeding sites was not affected by the areas, and the choice of this breeding site was due to the synanthropic habit of the mosquitoes sampled.

The variation in the degree of synanthropism revealed that some species have a greater preference for disturbed areas, which should be evaluated frequently, since mosquitoes previously considered wild have modified their habits and this could happen again.

The presence of several species with the potential to carry etiological agents found in the areas highlights the importance of epidemiological surveillance, especially in urban and peri-urban fragments. Studies like the current one provides important information about the species and enable the development of new and effective strategies aimed at efficient monitoring of culicidae in anthropic areas, contributing to public policies for urban expansion.

Mosquitoes draw the attention of authorities because they have been a public health issue for many years (Rueda, 2008). The number of vector species reported by this study is relevant, considering that with increasing environmental change, these species can participate in epidemiological cycles and the resurgence of some diseases (Guedes & Navarro-Silva, 2014).

*Aedes aegypti* is an exotic species that transmits dengue, yellow fever, zika, and chikungunya viruses (Menezes, 2016). *Aedes albopictus* is also an exotic species in Brazil, being capable of transmitting at least 22 arboviruses in Asia (Gratz, 2004). In Brazil it can transmit the Zika virus and laboratory experiments also point to the dengue virus, yellow fever, and chikungunya. However, females of *Ae. albopictus* have never been found naturally infected with these last three viruses in Brazil (Menezes, 2016).

*Haemagogus leucocelaenus* is the vector of the wild yellow fever virus (Zequi et al., 2005) and needs to receive special attention in areas with the presence of primates *Culex saltanensis* is a vector for some species of plasmodia and a species of triatomine, all of which parasitize birds (Zequi & Lopes, 2012). *Aedes terrens*, *Cx. eduardoi,* and *Li. durhamii* have never been found infected with any type of pathogen and *Tx. theobaldi* has no epidemiological importance since adult individuals do not perform hematophagy (Medeiros-Souza et al., 2013).

The data indicated the need to monitor these areas due to the presence of vector species and wild species in more altered environments. This adaptive capacity to disturbed environments can generate alterations in the epidemiological conditions of infectious agents (Guedes, 2012). Knowledge about this group is far from complete and more information about changes in biology, activity times, oviposition behavior, and susceptibility to insecticides contributes to the creation of new strategies to control and combat vector species in these areas close to human populations (Guedes, 2012).

## Appendix 17

- PCA resulting from the ordering of breeding sites according to abiotic variables.

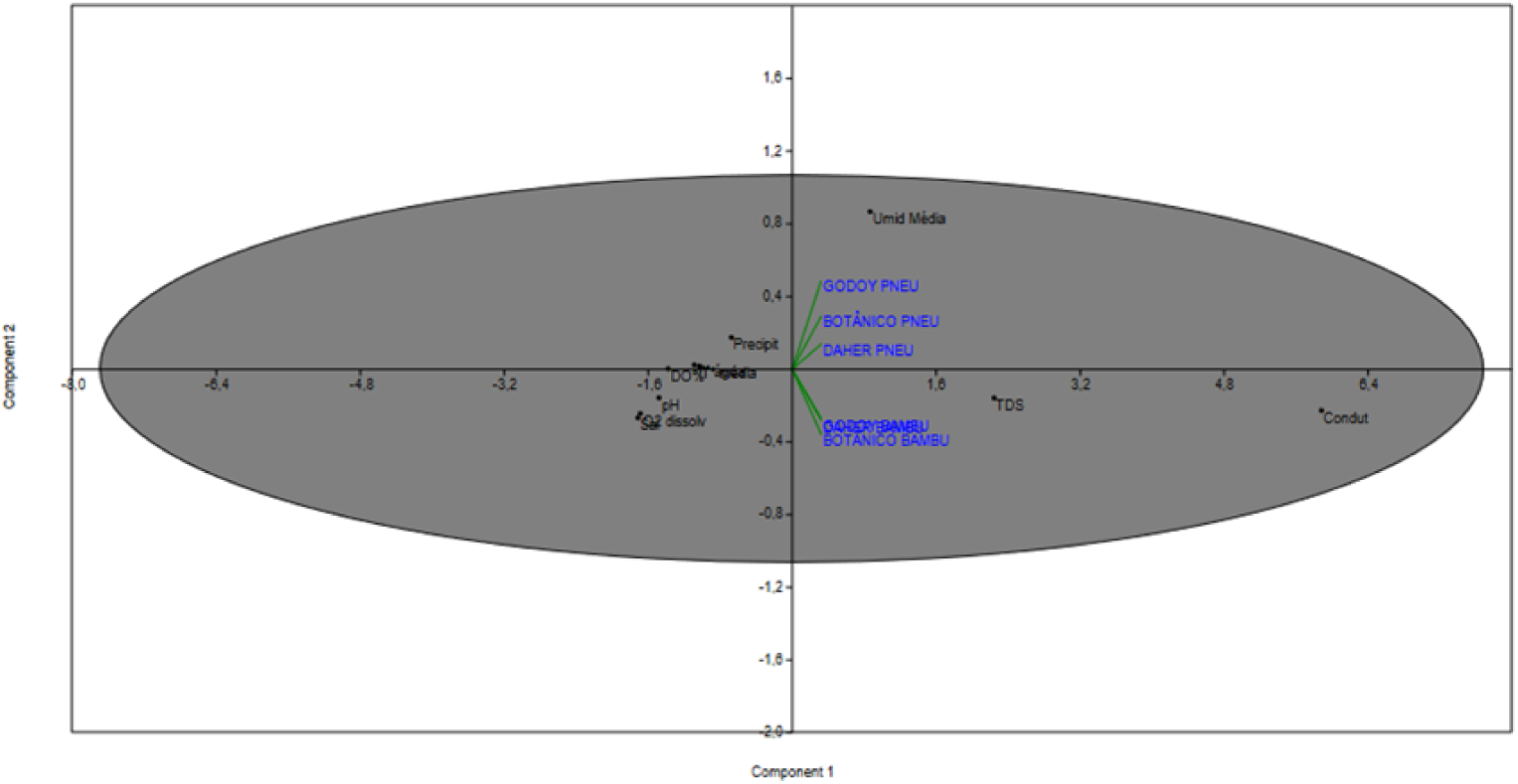

## Footnotes

1 Above sea level.

